# Single-trial Phase Entrainment of Theta Oscillations in Sensory Regions Predicts Human Associative Memory Performance

**DOI:** 10.1101/261487

**Authors:** Danying Wang, Andrew Clouter, Qiaoyu Chen, Kimron L. Shapiro, Simon Hanslmayr

**Author notes:** Address correspondence to Simon Hanslmayr.

## Abstract

Episodic memories are rich in sensory information and often contain integrated information from different sensory modalities. For instance, we can store memories of a recent concert with visual and auditory impressions being integrated in one episode. Theta oscillations have recently been implicated in playing a causal role synchronizing and effectively binding the different modalities together in memory. However, an open question is whether momentary fluctuations in theta synchronization predict the *likelihood* of associative memory formation for multisensory events. To address this question we presented movies and sounds with their luminance and volume modulated at theta (4 Hz), with a phase offset at 0° or 180° with respect to each other. This allowed us to entrain the visual and auditory cortex in a synchronous (0°) or asynchronous manner (180°). Participants were asked to remember the association between a movie and a sound while having their EEG activity recorded. Associative memory performance was significantly enhanced in the synchronous (0°) compared to the asynchronous (180°) condition. Source-level analysis demonstrated that the physical stimuli effectively entrained their respective cortical areas with a corresponding phase offset. Importantly, the strength of entrainment during encoding correlated with the efficacy of associative memory such that small phase differences between visual and auditory cortex predicted a high likelihood of correct retrieval in a later recall test. These findings suggest that theta oscillations serve a specific function in the episodic memory system: Binding the contents of different modalities into coherent memory episodes.

**Significance Statement:** How multi-sensory experiences are bound to form a coherent episodic memory representation is one of the fundamental questions in human episodic memory research. Evidence from animal literature suggests that the relative timing between an input and theta oscillations in the hippocampus is crucial for memory formation. We precisely controlled the timing between visual and auditory stimuli and the neural oscillations at 4 Hz using a multisensory entrainment paradigm. Human associative memory formation depends on coincident timing between sensory streams processed by the corresponding brain regions. We provide evidence for a significant role of relative timing of neural theta activity in human episodic memory on a single trial level, which reveals a crucial mechanism underlying human episodic memory.

## Introduction

A week after seeing your favorite band in concert, whilst sitting in your office, you mentally travel back and re-experience the band playing your favorite song. Episodic memory is the machinery that allows us to form, hold and revisit such rich, often multisensory, memories (Tulving, 2002). Yet sensory information is processed in specialized brain regions distributed across the cortex, which raises the fundamental question of how the different sensory elements are bound into a coherent memory representation and, more importantly, what affects the likelihood of that association being made. In this study, we address this question using a recently developed multisensory entrainment paradigm (Clouter, Shapiro, & Hanslmayr, 2017). We show that memory formation for dynamic audio-visual events can be predicted by momentary fluctuations in theta phase synchronization between sensory regions.

The hippocampus plays a fundamental role in episodic memory formation (Scoville & Milner, 1957) and receives highly preprocessed information from virtually all sensory modalities (Moscovitch, 2008; Muzzio, Kentros, & Kandel, 2009). This makes it a likely region supporting the binding function in memory. Importantly, within the hippocampus synapses are more likely to undergo modification if the inputs are active at the same time (Bliss & Collingridge, 1993). Theta oscillations – a dominant signal in the hippocampus (Jacobs, 2013) – synchronize neural ensembles (Buzsáki, 2010) and thus likely establish the synchrony required to effectively induce synaptic plasticity. Studies in rodents suggest that synaptic plasticity depends on the phase of the ongoing theta rhythm. Stimulation in the hippocampus yields either long-term potentiation (LTP) or long-term depression (LTD), depending on whether stimulation is applied at the peak or trough of the theta phase, respectively (Hölscher, Anwyl, & Rowan, 1997; Huerta & Lisman, 1995; Hyman, Wyble, Goyal, Rossi, & Hasselmo, 2003; Pavlides, Greenstein, Grudman, & Winson, 1988).This theta-phase-dependent learning mechanism has sparked considerable interest in human memory research and is an important component of several computational models of human memory (Hasselmo, 2005; Norman, Newman, Detre, & Polyn, 2006; Parish, Hanslmayr, & Bowman, 2017). However, direct evidence for such a theta-dependent synchronization mechanism in human memory is scarce. Such direct evidence would have to show two things: (i) externally synchronizing/de-synchronizing sensory regions in theta frequency impacts memory formation and (ii) momentary fluctuations in inter-areal phase synchronization predict memory formation. In the current study we investigate both predictions, going beyond a recent report wherein we addressed only the first of these hypotheses (Clouter et al., 2017).

## Materials and Methods

### Participants

Thirty-one healthy English-speaking young adults participated in the experiment (15 male; mean age: 21.48 years; range: 18 – 29 years). All participants had normal or corrected-to-normal vision and normal hearing. One participant was ambidextrous; all other participants were right-handed. Four participants were given course credits via the University of Birmingham’s Psychology Research Participation Scheme. The remaining 27 participants were paid £6 per hour for their participation. The data from three participants were excluded because of their poor, chance-level memory performance. The data from one participant was excluded due to a technical problem during EEG recording. The data from another three participants were excluded due to poor EEG data quality. The data from the remaining 24 participants were retained for the final data analysis.

### Stimulus Material

The visual and auditory stimuli used were taken from the same pool as those used in Experiment 3 of Clouter et al. (2017). Movie clips (N=192) were presented for three seconds in total 76 frames at a frame rate of 25 frames/s. The luminance of the movie clips was modulated from 0% to 100% with at 4 Hz with sine wave, with all movie clips initially starting at 50% luminance. On each trial, concurrent with the presentation of the movie, one of 192 sound clips was presented for three seconds, with a lag of 40 ms (i.e. starting 40 ms after the start of the movie). This 40 ms lag was necessary to account for the fact that auditory stimuli are processed faster than visual stimuli, and therefore to ensure a lag at the targeted phase difference of 0 and 180 between auditory and visual cortex (Clouter et al., 2017). The amplitude of the sound clips was modulated from 0% to 100% at 4 Hz with a sine wave, with all sound clips starting at 0% amplitude, and ramping in linearly. Each sound was modulated both at 0° and 180° phase offsets from sine-wave-modulated movies (accounting for the 40ms lag for the auditory stimuli). Each sound clip was from one of the eight sound categories: acoustic guitar, choir, movie soundtracks (two), electric piano, electric guitar, synthesizer, and an orchestra. Each sound category consisted of 24 sound clips. Each sound within a sound category was randomly assigned to a phase offset modulation with the constraint that the number of sounds for each phase offset modulation was equal. Each movie clip was randomly paired with a sound. The random assignment of phase offset modulation to a sound and a movie to a sound was done before the experiment and consistent across all the participants.

The experimental apparatus and stimulus presentation were identical to that used in Experiment 3 of Clouter et al. (2017). The experiment was programmed with MATLAB (R2013a; The Mathworks, Inc., Natick, MA, USA) using the Psychophysics Toolbox extensions (Brainard, 1997; Kleiner et al., 2007; Pelli, 1997). Presentation of visual stimuli were on a 21-inch CRT display with an nVidia Quadro K600 graphics card (875 MHz graphics clock, 1024 MB dedicated graphics memory; Nvidia, Santa Clara, CA, USA). Participants sat approximately 60 cm from the center of the monitor. The screen refresh rate of the monitor was 75 Hz. Auditory stimuli were presented with insert earphones (ER-3C; Etymotic Research, Elk Grove Village, IL).

### Experimental Procedures

Following the provision of informed consent, participants were prepared for EEG data collection, and were given instructions for, and familiarized with the tasks. During the formal experiment, participants were monitored by a web camera connected to a monitor in an adjoining EEG control room.

The experiment consisted of 12 blocks of an associative memory task, followed by synchrony judgment and unimodal source localizer tasks (see below). Each associative memory task block comprised an encoding phase, a distractor phase and an associative memory recall test phase. On each trial during the encoding phase, participants were presented with one of 16 movies along with one of 16 sound clips. Each trial started with a fixation cross with an inter-trial-interval between one and three seconds. Then, a sound-movie pair was presented for three seconds. After the presentation, participants were instructed to make a judgement as to how well the sound suited the contents of the movie by pressing the number keys on a keyboard. The instruction screen was presented until a response was made. The ratings ranged from 1 (the sound does not suit the movie at all) to 5 (the sound suits the movie very well). Participants were instructed to remember the association between the sound and the movie within each trial, upon which their memory would later be tested. Within each block, four sounds from four categories were associated randomly with the 16 movies with the constraint that the number of sounds for each phase offset condition was equal (i.e. two 0° phase offset modulated sounds and two 180° phase offset modulated sounds in one sound category).

The distractor phase was presented after the last trial of the encoding phase. Participants were presented with a random number on the screen, drawn randomly from 170-199. Participants were instructed to count backwards, aloud, by 3 starting from this number, for 30 seconds.

Participants were prompted on the screen to stop the distractor phase and start the associative memory recall test phase. In each block, the associative memory recall test phase consisted of 16 test trials, thus testing all of the 16 associations learned during the encoding phase. Each trial started with a fixation cross with an inter-trial-interval between one and three seconds. Participants were presented with one of the 16 sounds heard during the encoding phase for three seconds, along with four still images from four of the movies from the encoding phase. Then, participants were instructed to select the movie that was presented with the sound in the encoding phase using the number keys 1 through 4. The instruction screen was presented until a response was made. The movies from which to choose were, in the encoding phase, all presented with a sound from the same sound category.

The block presentation order was counterbalanced to ensure that each block was presented in each serial position an equal number of times across participants. Participants were allowed to take a break, if required, following each block. After the completion of the 12 associative memory task blocks, participants were given additional instructions for a synchrony judgment task. In this task participants were presented with 24 sound-movie pairs, drawn from the 192 sound-movie pairs. During the synchrony judgment task subjects were asked whether they could detect the synchrony (in-phase, 0° phase offset) or asynchrony (out-of-phase, 180° phase offset) between the flicker of the movie and sound. Participants were instructed to indicate asynchrony pressing number key 1 and indicate synchrony pressing 2.

Last, two unimodal source localizer tasks were run. Each consisted of 50 trials of flickered sound clips only, or 50 trials of flickered movie clips only. The sound and movie clips were drawn from the 192 sounds and movies used throughout the experiment. On each trial, participants were asked to rate how pleasant the sound or movie was by using the number keys 1 (the sound or the movie was very unpleasant) through 5 (the sound or the movie was very pleasant).

### EEG Recordings and Preprocessing

EEG was recorded from 128 scalp channels using a BioSemi ActiveTwo system. Vertical eye movements were recorded from an additional electrode placed 1 cm below the left eye. Horizontal eye movements were recorded from two additional electrodes placed 1 cm to the left of the left eye and to the right of the right eye. Online EEG signals were sampled at 1024 Hz by using the BioSemi ActiView software. The positions of each participant’s electrodes were tracked using a Polhemus FASTRAK device (Colchester, Vermont, USA) and recorded by Brainstorm (Tadel, Baillet, Mosher, Pantazis, & Leahy, 2011) implemented in MATLAB.

Offline EEG data was preprocessed using the Fieldtrip toolbox for MEG and EEG analysis (Oostenveld, Fries, Maris, & Schoffelen, 2011). The continuous EEG data was bandpass filtered between 1 and 100 Hz and bandstop filtered between 48 – 52 Hz and 98 – 102 Hz to remove potential line noise at 50 Hz and 100 Hz. The data were then epoched from 2000 ms before stimulus onset to 5000 ms after stimulus onset, and downsampled to 512 Hz. Bad channels and trials with coarse artefacts were excluded by visual inspection before applying an independent component analysis (ICA). In the unimodal conditions, one bad channel was excluded for each of five participants. Four bad channels were excluded for another one participant. In the multimodal conditions, one bad channel was excluded for each of three participants. Two bad channels were excluded for another one participant. Four bad channels were excluded for each of another two participants. ICA components suggesting eye movement artefacts and regular pulse artefacts were removed from the data. Bad channels were interpolated by the method of triangulation of nearest neighbors based on the individuals’ electrode positions. After re-referencing the data to average, trials with artefacts were manually rejected by visual inspection. Participants with fewer than 16 artefact-free trials in any of the conditions of interest were excluded from further analysis. The mean number of trials remaining in each condition were: unimodal sound: 42 (25 – 50), unimodal movie: 42 (26 – 49), multimodal 0 remembered: 34 (21 – 48), multimodal 0 forgotten: 43 (32 – 55), multimodal 180 remembered: 31 (22 – 44), multimodal 180 forgotten: 46 (28 – 61).

For three participants who provided their own MRI scans, their head models were created based on their structural scans (cf. (Michelmann, Bowman, & Hanslmayr, 2016)The MRI scans were segmented into four layers, brain, cerebrospinal fluid, skull and scalp by using the Statistical Parametric Mapping 8 (SPM8) toolbox (http://www.fil.ion.ucl.ac.uk/spm) and the Huang toolbox (Huang et al., 2013). Then a volume conduction model was constructed by calling the ‘dipoli’ method implemented in the Fieldtrip toolbox. Individuals’ electrode positions were aligned to their head models. For further group analysis across all 24 participants, each individual MRI was warped to an MNI template MRI provided by the Fieldtrip toolbox then the inverse of the warp was applied to a template dipole grid to have a same location of each grid point for each participant in normalized MNI space. For the remaining participants who did not have their own MRI scans, the MNI template MRI and a template volume conduction model were used. Individuals’ electrode positions were aligned to the template head model. Source models were prepared with the template volume conduction model and the aligned individuals’ electrode positions.

### Unimodal Source Localization

In the unimodal sound condition, EEG data from the unimodal sound condition was scalp current density (SCD) transformed using the finite-difference method (Huiskamp, 1991; Oostendorp & Oosterom, 1996). In addition, the leadfields that were computed based on scalp potentials were SCD transformed by applying the transformation matrix that was used for transforming the potential data. This step is necessary for source localization and reconstruction of data from correlated with brain sources (such as data from auditory sources) using the beamforming method (Murzin, Fuchs, & Scott Kelso, 2013).

Source activity was reconstructed using a linearly constrained minimum variance (LCMV) beamforming method (Van Veen, Van Drongelen, Yuchtman, & Suzuki, 1997). In the movie condition, source analysis was calculated for individual electrode positions, individual volume conduction model and normalized grid positions for the three participants who had their MRI scans. For the other participants, source analysis was run with individual electrode positions, grid positions and template volume conduction model. All the analyses were run on potential (i.e. average referenced) data. In the sound condition, source analysis was conducted using SCD-transformed leadfields on SCD-transformed data. Time series data was reconstructed on 2020 virtual electrodes for each participant. Event-related potential (ERP) was calculated at each virtual electrode for each unimodal condition. Time-frequency analysis was applied to each source ERP with a Morlet wavelet (width = 7) using a frequency of interest between 3 and 5 Hz. Evoked power was averaged between 0.75 and 2.75 seconds post stimulus onset and between 3.5 and 4.5 Hz. A baseline condition was created by randomly assigning each trial to 0°, 90°, 180°, or 270° phase offset by moving the signal onset forward in time by 0, 32, 64, or 96 samples (0, 62.5, 125, or 187.5 ms) with a restriction that the number of trials in each phase offset were approximately equal. The evoked power of the baseline condition was therefore expected to be minimal at corresponding auditory and visual sources. The evoked power differences were grand averaged across participants and the grand average evoked power differences were interpolated to the MNI MRI template. The coordinates for auditory and visual regions of interest (ROIs) were determined by where the maximum grand average evoked power differences were shown.

### Multimodal Source Reconstruction

To reconstruct the time series data from auditory sources, the multimodal data was SCD-transformed. Beamforming analysis has a notoriously poor performance in reconstructing highly correlated source activity such as during auditory stimulation. A solution for EEG source localization was proposed by Murzin et al. (2013) who suggest that source analysis should be computed based on the scalp electrodes over left and right hemispheres separately(Murzin et al., 2013). As a result, the spatial filter that was computed with only left scalp electrodes was applied to the left scalp time series data and the spatial filter that was computed with only right scalp electrodes was applied to the right scalp time series data. The time series data therefore corresponded to the sources of the left hemisphere and right hemisphere, respectively. The time series data at the left auditory ROI was extracted at the pre-determined left auditory coordinate from the unimodal source localization step. Similarly, data at the right auditory ROI was extracted at the pre-determined right auditory coordinate. The signals were then averaged across left and right auditory ROIs. The time series data at visual ROI was reconstructed without SCD transformation and was extracted at the pre-determined visual coordinate.

To solve the problem of random direction of source dipoles caused by the LCMV beamformer, each participant’s left and right auditory source ERPs were plotted along with visual source ERPs in each phase offset condition. The sign of the source reconstructed time series data was flipped in direction if any source ERPs showed the opposite of the expected direction of the visual and auditory ERP components (P1, N1 or P2) by multiplying the data in the time series by −1. The same approach was taken if the time series showed the opposite of the expected entrainment (with respect to the flicker of the physical stimuli) (e.g. if the 0° phase offset condition looked like 180°, or the 180° phase offset condition looked like 0°). This flipping procedure was applied consistently across all trials, regardless of condition. A control analysis was conducted to confirm that the phase relationships between 0° and 180° conditions in the auditory and visual ROIs remained constant between before and after flipping. All distributions were non-uniform as indicated by the Rayleigh test. Thus the flipping procedure only resulted in a better illustration of the entrainment effect but did not bias the results in favor of our hypothesis.

### Statistical Analysis

To compute the instantaneous phase differences, the source reconstructed time series data were bandpass filtered between 1.5 and 9 Hz. Then, the ERPs were computed for each phase offset condition at each source. The Hilbert transformation was applied to the source grand averaged ERPs that were redefined in time (1000 ms before stimulus onset to 4000 ms after stimulus onset). The instantaneous phases were calculated from the Hilbert transformed data and unwrapped. The instantaneous phase differences were calculated between auditory and visual sources for one second, beginning one second after stimulus onset (to avoid influences of onset and offset responses) in each phase offset condition. The Rayleigh test and V test were used to test circular uniformity of the instantaneous phase differences in each phase offset condition.

For the subsequent memory analysis, the Hilbert transformation was applied to the filtered single-trial source data. The instantaneous phases of each trial were calculated from the Hilbert transformed data and unwrapped. The instantaneous phase differences were calculated between auditory and visual sources for one second, beginning one second after stimulus onset. The single-trial phase entrainment measure in each phase offset condition was computed for each trial and each time point by calculating the resultant vector length of a vector that consisted of the data value at the time point and the theoretical phase offset (0° or 180°). The resultant vector length was collapsed across time between 1 and 2 seconds, resulting in one value per trial. A 2 (phase offset condition) x 2 (subsequent memory) repeated-measures ANOVA was conducted on the averaged single-trial phase entrainment values for remembered and forgotten items in each phase offset condition.

To further test whether the degree of single-trial phase entrainment predicts subsequent memory performance, a logistic regression model was fit using the single-trial phase entrainment measure as the predictor, and memory performance on each trial as the dependent variable, resulting in model slopes for each phase offset condition. These slopes were normalized for each participant. The normalized slopes were then averaged between the two phase offset conditions for each participant. A one sample t-test was conducted to test whether the slopes were significantly larger than 0.

To get a more fine-tuned picture of the relationship between single-trial phase angle and memory performance, we calculated the mean phase direction of the time points between 1 and 2 seconds for each trial. Depending on the mean phase direction of a trial, we sorted the trials into four bins, centered at 0°, 90°, 180° and 270° each with a bin width of ±45°. To reduce the trial-number bias in each bin, we randomly selected 12 trials (the minimum trial number was 13) from each bin and calculated the proportion of remembered trials out of the 12 trials in each bin. This procedure was repeated 10 times and averages across these 10 repetitions were built for each subject to reflect the proportion of remembered trials in each phase bin. A repeated-measures ANOVA was conducted on the memory accuracy across the four bins.

## Results

### Behavioral performance

We replicated the previous findings (Clouter et al., 2017) by showing that memory performance in the 0° phase offset condition was significantly better than in the 180° phase offset condition (Figure 2) using a paired-samples t-test, *t* (23) = 2.069, *p* = 0.025 (one-sided). To rule out the influence of perceptual factors on the memory effect, a sensitivity index (*d’*) was computed for perceptual judgment performance on synchronous and asynchronous stimuli. One-sample t-tests showed that the *d’* for participants’ judgment on synchronous stimuli vs. asynchronous stimuli did not differ from zero (synchronous, *t* (23) = 1.317, *p* = 0.201; asynchronous, *t* (23) = 1.008, *p* = 0.324). In addition, a paired-samples t-test showed that *d’* for discriminating synchronous from asynchronous stimuli did not differ from *d’* for discriminating asynchronous stimuli from synchronous stimuli, *t* (23) = 0.074, *p* = 0.942. These results suggest that participants were unable to discriminate synchronous stimuli from asynchronous stimuli, thus ruling out the contribution of basic perceptual confounds to the observed memory effect.

**Figure 1.**
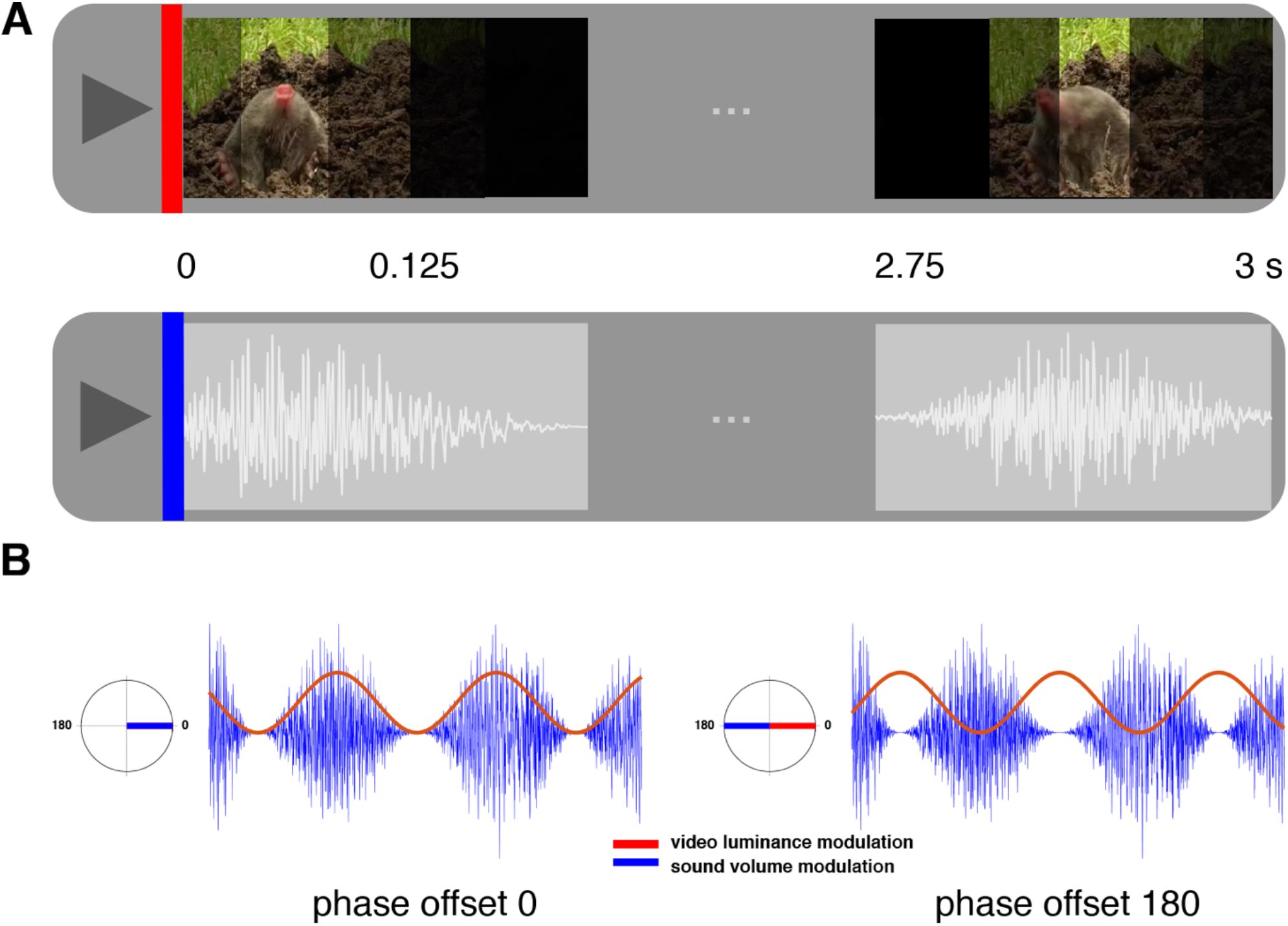
Stimulus material of associative memory encoding task. ***A***, An example of visual and auditory stimuli presented in the associative memory encoding task. Top: Movies were luminance modulated with a sine wave, and so flickered visually at 4 Hz. Bottom: Sounds were amplitude modulated with a sine wave, and “flickered” at 4 Hz. The movie and the sound are flickering in phase (i.e., with a 0° phase offset). ***B***, Sine-wave-modulated visual (red) and auditory (blue) stimuli. Left: the sound clip was amplitude modulated at 0° phase offset from the sine wave that modulated the luminance of the movie. Right: the sound clip was amplitude modulated at 180° phase offset from the since wave that modulated the luminance of the movie. Note, that the auditory stimulus always lagged 40ms with respect to the visual stimulus, which is not shown here for illustration purposes.

**Figure 2.**
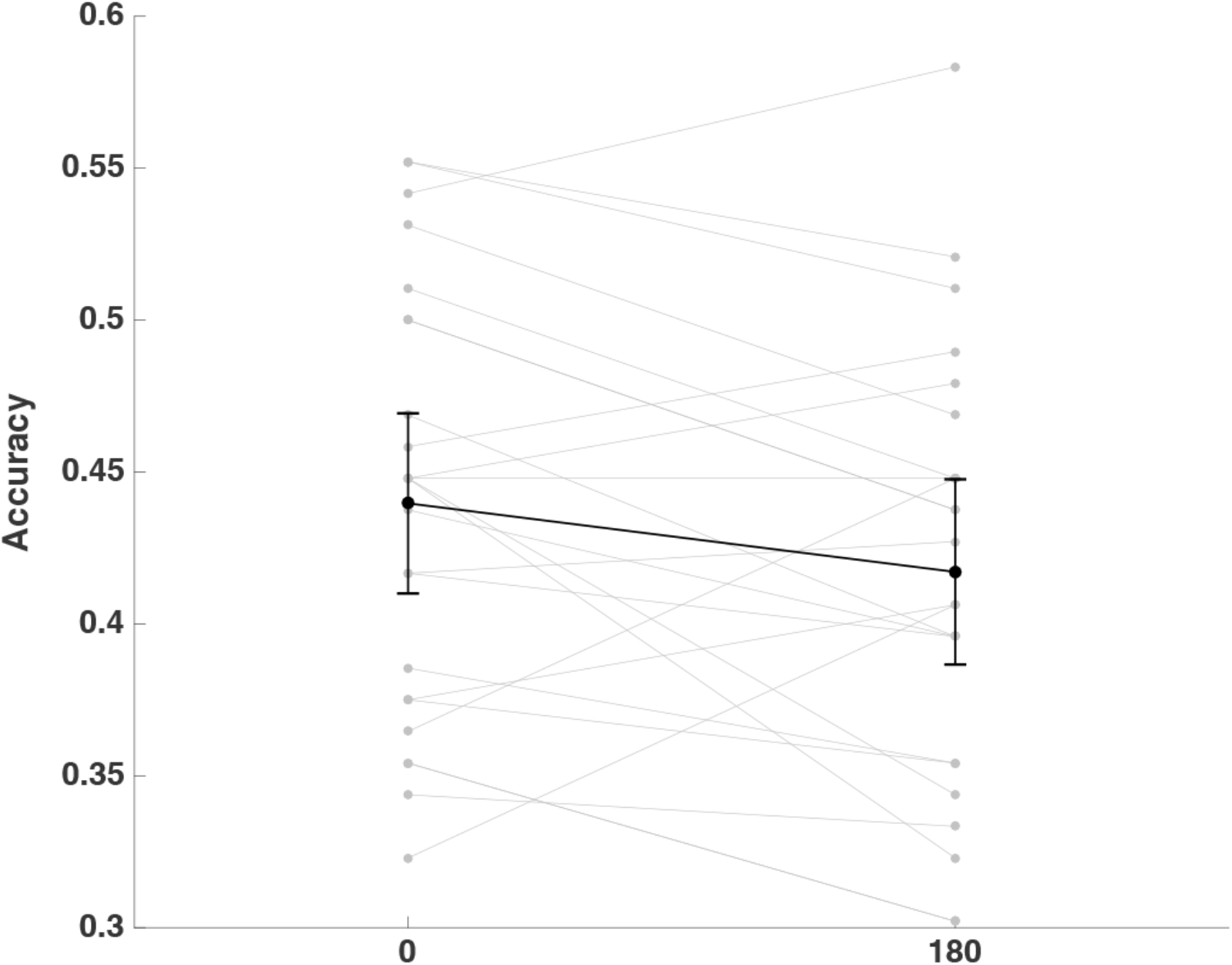
Associative memory task performance. Proportion of the correctly selected movie scenes that were associated with presented sounds in each phase offset condition. Note that chance level is at 25%. Error bars represent 95% confidence intervals of means in 0 and 180 phase offset conditions. Individual data for correct associative memory performance in 0 and 180 phase offset conditions is shown in grey.

To investigate if participants’ subjective judgment on how well a sound suited the contents of a movie influenced memory performance, a 2 (subsequent memory) x 2 (phase offset condition) repeated-measures ANOVA was performed on the average rating score. A significant main effect of subsequent memory was found (mean rating score in subsequently remembered items: 3.038, mean rating score in subsequently forgotten items: 2.827, *F* (1, 23) = 24.224, *p* < 0.001). A significant main effect was also shown for phase offset, mean rating score in 0° phase offset condition, 2.979, mean rating score in 180° phase offset condition, 2.886, *F* (1, 23) = 7.962, *p* = 0.010. These results suggest that the observed better memory performance for 0° phase offset condition compared to 180° condition could be driven by perceived semantic congruency between multi-sensory stimuli. To control for this possible alternative, we conducted an analysis where the trial numbers in each rating scale were equalized between the 0° phase offset condition and the 180° phase offset condition by randomly sub-selecting trials for each participant (mean trial number 170, range 148 - 190). Associative memory accuracy was calculated based on these selected trials – now equalized for semantic congruency – for each participant. This procedure was repeated 10 times. The resultant associative memory accuracy was averaged for each participant across 10 repetitions. The memory advantage in the 0° phase offset condition was still valid, as supported by a paired-samples t-test, *t* (23) = 1.903, *p* = 0.035 (one-sided), thus confirming our original results.

### Phase in sensory cortices is entrained to multi-sensory stimulation

The ROIs obtained from the unimodal source localization results are shown in Figure 3. Multimodal grand average ERP waveforms were extracted from the visual and auditory ROIs determined by these unimodal source localization results. Figure 4 shows the grand average waveforms for the 0° and 180° phase-offset conditions for the visual and auditory ROIs. Rayleigh tests and V tests were conducted for the 0° and 180° phase offset conditions separately, using the data points of instantaneous phase differences between auditory and visual signals from 1 to 2 seconds (n = 513 samples) as dependent variable. Both tests enable rejection of the null hypothesis that the distribution of the phase differences in the 0° phase offset condition or the 180° phase offset condition are uniformly distributed; the resultant vector length in the 0° condition: 0.9197, *p* ≈ 0; resultant vector length in 180° condition: 0.9192, *p* ≈ 0. The V test further suggests that the distribution of phase differences in each phase offset condition has a mean direction of its entrained phase, 0° or 180° (both *p*s = 0), respectively. These results suggest that our paradigm was effective in synchronizing (0° condition) or desynchronizing (180° condition) the auditory and visual cortex in the theta frequency, replicating Clouter et al. (2017).

**Figure 3.**
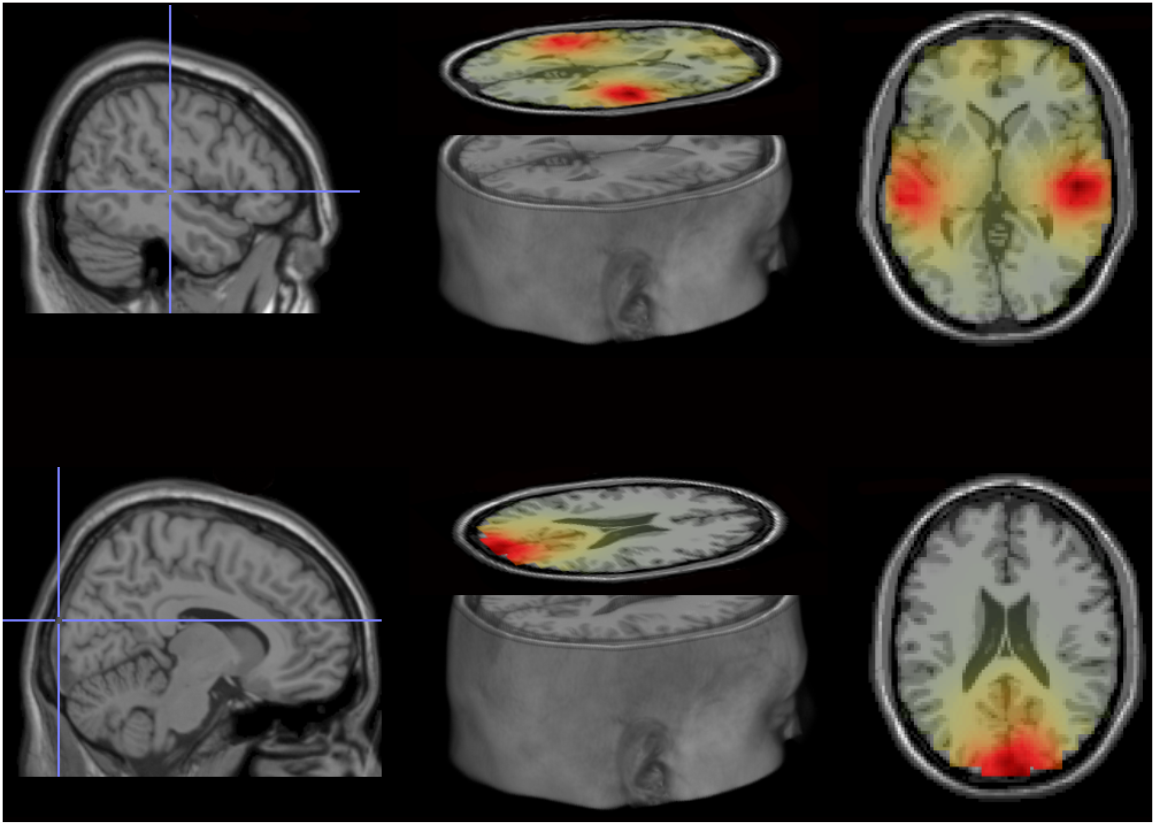
Source localization of theta power in the unimodal conditions. Top: Auditory sources, MNI coordinates of ROIs: right, 50, −21, 0; left, −60, −29, 0. Bottom: Visual source, MNI coordinates of ROI: 10, −99, 20. Evoked power was averaged over 3.5 and 4.5 Hz, between 0.75 and 2.75 s at each virtual electrode in the unimodal movie and sound conditions and the baseline conditions (see Materials and Methods). Grand average power differences between unimodal conditions and baseline conditions were interpolated to a MNI MRI template. The source coordinates were determined by where the maximum grand average power differences were. The source plots were generated by the Fieldtrip toolbox and MRIcro for OSX (http://www.mccauslandcenter.sc.edu/crnl/mricro).

**Figure 4.**
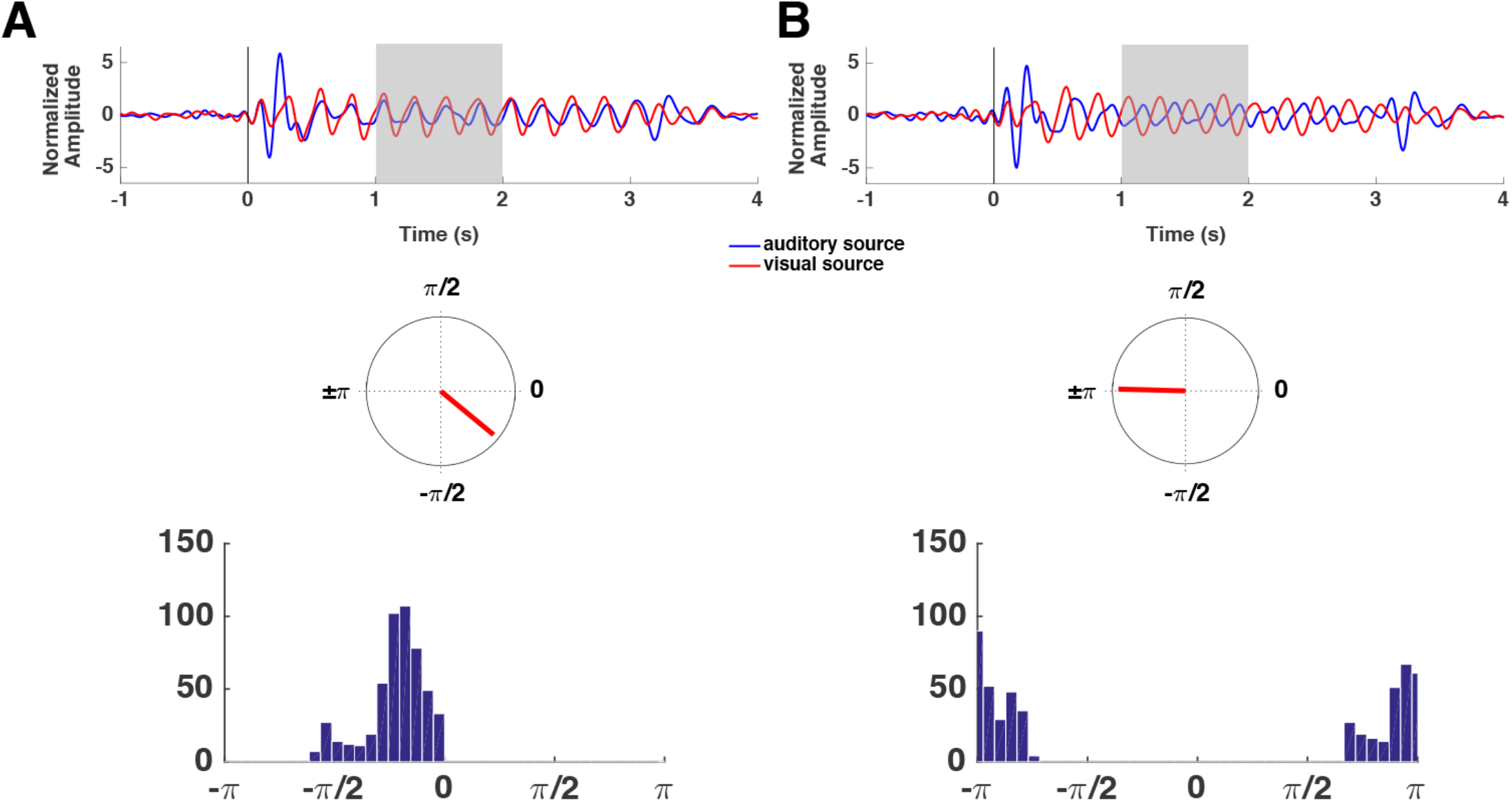
Phase differences between auditory and visual sources in each phase offset condition. ***A***, Phase differences between auditory and visual sources in 0° offset condition. Top: Amplitude normalized grand average ERP signals at auditory (blue) and visual sources (red). Middle: Mean resultant vector of instantaneous phase differences between auditory and visual sources, between 1 and 2 s (shaded time window on the grand average ERPs) is plotted on a unit circle. Bottom: Histogram of wrapped instantaneous phase differences between auditory and visual sources, between 1 and 2 s, using 40 equally-sized bins. ***B***, As in A, but for the 180° offset condition.

### Single trial phase entrainment predicts subsequent memory performance

To investigate our main hypothesis, i.e. whether phase differences between auditory and visual regions predict the likelihood of memory formation, a single-trial phase entrainment measure was calculated for each phase offset condition (see Materials and Methods). To this end the resultant vector length for every time point was calculated between the observed phase difference (between the auditory and visual cortices) and the phase difference of the physical stimuli (i.e. either 0° or 180°). This measure yields values close to 1 for high entrainment, where a given trial closely followed the rhythmic stimulation, and values close to 0 for low entrainment, where a given trial deviated from the rhythmic stimulation. Figure 5 illustrates this measure with an example from four single trials. Figure 5A and 5B shows single trials from 0° offset condition. The trial for a later remembered stimulus reveals a strong entrainment at 0°, while the forgotten trial shows a small value. In contrast, the forgotten trial in the 180° offset condition shows a strong entrainment to 180°, whereas the remembered trial shows a weak entrainment to 180° (Figure 5C and 5D). This pattern is consistent with the notion that 0° is optimal and 180° is non-optimal for the formation of associative memories. To test this idea formally, we conducted a 2 x 2 ANOVA with the factors phase offset condition (0° vs. 180°) and subsequent memory (hits vs misses), expecting to find a significant interaction. Indeed, such a significant interaction between phase offset condition and subsequent memory was observed, *F* (1, 23) = 4.627, *p* = 0.042. Subsidiary paired-samples t-tests showed that this interaction was the result of a trend for stronger phase entrainment in remembered trials than forgotten trials in the 0° phase offset condition, *t* (23) = 1.496, *p* = 0.074 (one-sided), and significantly weaker entrainment in remembered trials than in forgotten trials in the 180° phase offset condition, *t* (23) = −2.165, *p* = 0.021 (one-sided) (Figure 6A). Together, these results reveal that the likelihood of a movie and a sound being successfully associated in memory varies with the strength of entrainment.

**Figure 5.**
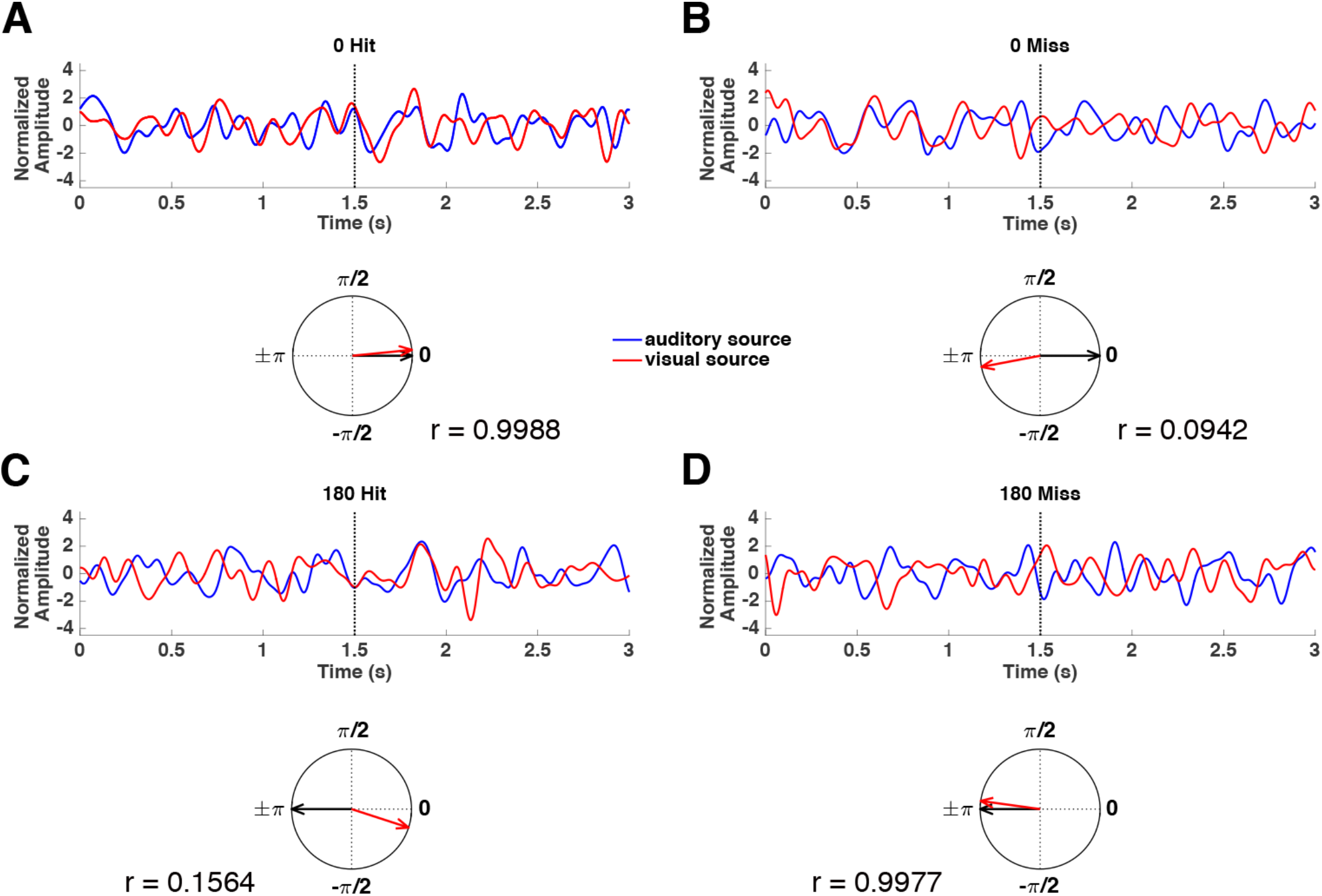
An example of four single trials in 0° remembered, forgotten conditions and 180° remembered, forgotten conditions, respectively. ***A***, Top: A band-pass filtered (1.5-9 Hz) single trial from auditory (blue) and visual (red) sources in 0° offset remembered condition. Amplitude was normalized. Bottom: instantaneous phase difference between auditory and visual sources at a certain time point 1.5 s of the single trial is plotted on a unit circle (red line). Single trial phase entrainment was calculated as the resultant vector length between the measured phase difference (red arrow) and the entrained phase offset 0° (black arrow), which was 0.9988 in this case. ***B***, same as in A but for 0° offset forgotten; ***C***, same as in A but for a 180° offset remembered trial; ***D***, same as in A but for a 180° offset forgotten trial.

**Figure 6.**
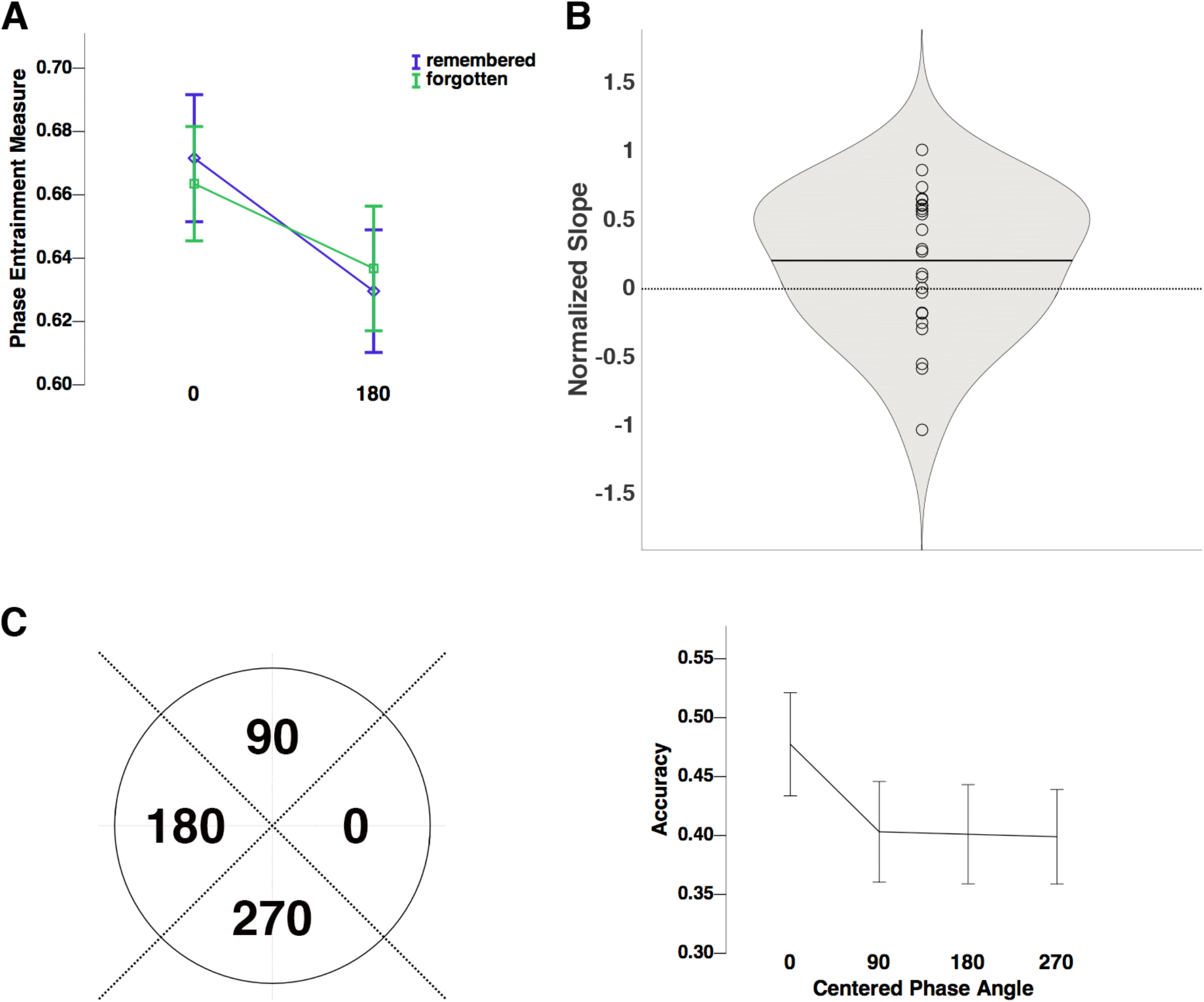
Memory performance as a function of phase entrainment. ***A***, Phase entrainment measure was averaged between 1 and 2 s for each trial. These single-trial phase entrainment values are plotted as a function of subsequent memory (remembered: purple; forgotten: green) in each phase offset condition. ***B***, Normalized slopes of logistic regression models for each phase offset condition were averaged between conditions for each subject. The logistic regression model was fitted to each subject’s memory performance, against the single-trial phase entrainment measure (averaged between 1 and 2 s). The violin plot represents the distribution of the averaged normalized slope values. The black horizontal bars represent the mean of the data. Individual scatter points represent the averaged normalized slope values for each participant. Violin plots were created with the MATLAB extension violin.m (Hoffmann, 2015). ***C***, Right: Four bins were centered at 0°, 90°, 180°, 270°, with a bin boundary of ±45°. Left: Recall accuracy was calculated in each bin. The mean phase direction between 1 and 2 s was computed for each trial. Each trial was sorted into one of the four bins depending on the mean direction. The proportion of remembered trials was calculated for 12 trials that were randomly selected from each bin for each iteration. The recall accuracy was averaged across 10 iterations.

Hits were associated with strong entrainment in the 0° phase offset condition but low entrainment in the 180° phase offset condition, whereas misses showed the opposite pattern.

Another way to describe this interaction is that the closer the audio-visual phase difference in a given trial is to 0, the higher the likelihood that the sound and the movie are associated in memory. To test this prediction more directly we measured the single-trial phase entrainment to 0° in both phase offset conditions. We then fitted a logistic regression model by using single-trial phase entrainment to 0° to predict memory outcome in each phase offset condition for each participant. The normalized slopes of each participant were averaged across two phase offset conditions. As shown in Figure 6B, one-sample t-tests revealed that the values of the slopes were significantly positive, *t* (23) = 1.958, *p* = 0.031 (one-sided). This result indicates that associative memory performance increases as a function of synchronization between the visual and auditory cortex, i.e. the closer their phase difference is to 0°.

To illustrate this relationship between audio-visual synchronization and memory performance in a more fine-grained manner, single trials were sorted into four bins centered at 0°, 90°, 180°, and 270°. To account for possible trial-number bias, we equalized the number of trials in each bin by randomly selecting 12 trials from each bin. Then the proportion of remembered trials was calculated for each bin with this procedure repeated 10 times for each participant. Finally, the proportion of remembered trials calculated at each iteration was averaged across those 10 iterations. The results show that the recall accuracy in the 0° bin was significantly higher than the accuracy in any of the other three bins (Figure 6C), which did not differ from each other. A repeated-measures ANOVA with the factor phase bin (0°, 90°, 180°, 270°) statistically confirmed this effect on recall accuracy, *F* (3, 69) = 6.014, *p* = 0.001. Pairwise t-tests showed that accuracy in the 0° bin was significantly higher than in the 90°, 180°, and 270° bins, *t* (23) = 3.061, *p* = 0.006; *t* (23) = 2.739, *p* = 0.012; *t* (23) = 4.293, *p* < 0.001. Whereas recall accuracy in the 90°, 180°, and 270° bins did not statistically differ from each other (all *p*s > 0.8). This result replicates the behavioral findings of Clouter et al. (2017), and suggests further that there is a narrow time window around 0 degrees for phases to synchronize in order to optimize the formation of associative memories.

## Discussion

We investigated a fundamental question in the field of memory research, i.e., does the degree of theta phase synchronization between visual and auditory cortical regions predict successful encoding of associations between visual and auditory information. We used a recently developed multisensory entrainment paradigm (Clouter et al., 2017) and replicated the findings from Clouter et al. (2017) by showing that episodic memory performance was significantly enhanced when the multisensory information was presented synchronously at 4 Hz. We used the physical sensory stimuli required to be associated to entrain the respective visual and auditory cortices at the intended phase offsets, successfully replicating Clouter et al. (2017). Importantly, however, our results go beyond the previous findings of Clouter et al. to reveal that the single trial phase synchrony between visual and auditory cortex *predicts* subsequent success on encoding the association between the sensory information into long-term memory. This finding is crucial because it reveals the underlying mechanism of synchronous presentation between sensory stimuli at theta frequency enhancing episodic memory. Moreover, the findings provide additional evidence as to the causal role of theta phase in human episodic memory formation by showing that synchronized theta activity between sensory information processing areas is a critical mechanism to bind multi-sensory information into episodic memory on a trial-by-trial basis.

Computational models suggest that the hippocampus binds diverse cortical representations into a sparse unified episodic memory representation by rapid synaptic modifications between cortical inputs and hippocampal neurons through LTP (McClelland, McNaughton, & O’Reilly, 1995). The induction of LTP depends on the coordinated timing of action potentials across populations of neurons as demonstrated by the animal literature *in vitro* and *in vivo* that stimulation at the peak of hippocampal theta phase induces LTP (Hölscher et al., 1997; Huerta & Lisman, 1995; Hyman et al., 2003; Pavlides et al., 1988). Given the causal role of theta phase in the induction of LTP, the relative timing between cortical inputs and hippocampal theta oscillations should also play a causal role in human episodic memory formation. However, most evidence is correlational and post-hoc (Backus, Schoffelen, Szebényi, Hanslmayr, & Doeller, 2016; Fell et al., 2003; Rutishauser, Ross, Mamelak, & Schuman, 2010). Clouter et al. (2017) provided the first causal evidence in humans by showing that associative memory performance was significantly better in the 0° condition than the 180° condition. The present study successfully replicates the findings from Clouter et al. (2017) although our effect is weaker and with more variability. One possible reason is that half of our stimuli was not used in Clouter et al. (2017), which might cause variability in entraining visual and auditory regions, hence reducing the effect. However, this variability allowed us to link trial-by-trial fluctuation in phase entrainment with subsequent memory performance. The strength of single trial entrainment in the 0° condition was positively related to episodic binding between visual and auditory information in that trial. In contrast, for the 180° condition stronger entrainment was related to a failure in memory formation. To put it another way, the closer the 4 Hz phase difference between the two sensory regions was to 0° the better the memory outcome (Figure 6B). This finding extends our previous results in that it directly links the theta phase dynamics with memory performance on a single trial level, suggesting that the relative timing in theta neural oscillations is a main driving force underlying long-term associative memory formation.

Learning of associations between unrelated elements is highly dependent on the hippocampus (Eichenbaum & Cohen, 2004; Gonzalo, Shallice, & Dolan, 2000; Staresina & Davachi, 2009). Although we were not able to observe signals from the hippocampus, the hippocampus is strongly implicated as the candidate region influencing the subsequent memory effect. Successful encoding of associations between cortical inputs happens when the inputs arrive at the peak of hippocampal theta phase triggering strong LTP in the synapses (Hasselmo, 2005). In human studies, functional connectivity between neocortical areas and the MTL has been related to successful episodic encoding (Schott et al., 2013; Summerfield et al., 2006). Moreover, item-context binding and integration between new information and old mnemonic trace has been suggested to be supported functionally by MTL theta power, as well as theta coupling between the MTL and cortical regions (Backus et al., 2016; Staudigl & Hanslmayr, 2013). Therefore, the current finding that synchronization between visual and auditory theta activity predicts later memory success could reflect that coincident volleys of action potentials terminating in the MTL at similar theta phases, which is then more likely to induce hippocampal LTP in the synapses of the respective auditory and visual downstream neural ensembles in the MTL.

One could argue that the observed phase synchronization between early sensory areas reflects multisensory integration, which captures attention and results in better memory (Senkowski, Schneider, Foxe, & Engel, 2008). However, the observed interaction between strength of entrainment and subsequent memory rules out the explanation that successful encoding of a sound-movie pair depends on early sensory level attention. The strength of entrainment to a periodic stimulus is positively related to attention, with attention increasing the entrainment response (Lakatos, Karmos, Mehta, Ulbert, & Schroeder, 2008; Müller et al., 2006; Nozaradan, Peretz, & Mouraux, 2012; Saupe, Schröger, Andersen, & Müller, 2009). If attention at the sensory level is the sole factor modulating successful memory formation, phase entrainment should be significantly stronger for remembered trials than forgotten trials, regardless of the phase offset condition (i.e. 0 vs 180), which is not what we observed. Furthermore, the interaction effect between subsequent memory and phase offset condition was shown only between 1 to 2 seconds after stimulus onset, not before or after. This time course suggests that the memory formation was not influenced by early evoked responses or stimulus-related phase coupling typically reported in the multisensory integration literature (Senkowski et al., 2008) but influenced by synchronized theta activity in a later and more sustained fashion, which is arguably a mechanism of successful encoding of associations (Summerfield & Mangels, 2005).

A remarkable finding is the fact that the degree of phase entrainment showed a reverse function of subsequent memory in the 180° condition. This raises the question of what the underlying mechanism of the weaker entrainment to 180° for remembered trials is. This perturbation from 180° points towards a resilience mechanism in the brain, preventing it from being entrained to a non-optimal phase offset in order to get closer to a more optimal phase for forming a memory association (i.e. 0°). Hippocampo-cortical loops are plausible candidates for such a mechanism, which fits with fMRI findings showing that increases in hippocampal activity are correlated with spatiotemporal desynchronization between elements that form a memory (Staresina & Davachi, 2009). Therefore, the hippocampus might put in more work when encoding associations between stimuli that are asynchronously presented. However, how the hippocampus feeds back the cortical information so that it can avoid being entrained remains unknown. A possible explanation could be that the overall hippocampal theta power increases when phase difference gets closer to 0° relative to when the inputs are strongly entrained to 180°, creates a ‘theta state’ conducive to episodic binding. If this is the case, memory should also slightly benefit from 90° and 270° phase offsets. However, when sorting single trials into four phase bins corresponding to 0°, 90°, 180° and 270° memory performance in the 0° condition was significantly better than in any of the other three conditions, with 90, 180 and 270 showing equally poor performance levels. This pattern perfectly replicates the pattern obtained by Clouter et al. (2017) and suggests that there is a narrow time window around <62.5 ms during which neural events have to coincide in order to result in effective memory formation. One such fine grained learning mechanism is Spike-Timing-Dependent-Plasticity (Song, Miller, & Abbott, 2000). However, whether STDP is involved in the observed effects needs to be further investigated.

In conclusion, our findings support the notion that externally induced inter-regional theta phase synchronization supports associative memory formation on a trial-by-trial basis. By successfully replicating findings from a previous study that used the same paradigm (Clouter et al., 2017), we further show that the degree of multisensory entrainment causes subsequent remembering and forgetting. Stronger entrainment in the 0° condition leads to successful memory while stronger entrainment in the 180° condition relates to failure in episodic binding. Such a finding supports that human episodic memory formation indeed depends on relative timing between the cortical processing of sensory streams, in turn providing a powerful tool to predict single trial memory formation based on the performance of entrainment on any given trial. With this experimental paradigm, neural synchronization can be precisely controlled to test how synchronized neural activity influences recollection vs. familiarity and monitor whether the current brain state is at encoding or retrieval (Ezzyat et al., 2017; Rizzuto, Madsen, Bromfield, Schulze-Bonhage, & Kahana, 2006), given the role of different hippocampal theta phases in episodic memory processes (Hasselmo, 2005; Hasselmo, Bodelón, & Wyble, 2002). Clinically, the paradigm could provide a non-invasive and accessible way to enhance episodic memory performance via external synchronization in healthy subjects and patients suffering from memory disorders such as Alzheimer (Huerta & Lisman, 1995; Iaccarino et al., 2016).

## Acknowledgements

This research was funded by the ERC grant Code4Memory (Grant Agreement 647954) awarded to SH, who is further supported by the Wolfson Society and the Royal Society.

